# Circadian rhythm in cortical chloride homeostasis underpins variation in network excitability

**DOI:** 10.1101/2021.05.12.443725

**Authors:** Enrico Pracucci, Robert Graham, Laura Alberio, Gabriele Nardi, Olga Cozzolino, Vinoshene Pillai, Luciano Saieva, Darren Walsh, Silvia Landi, Jinwei Zhang, Andrew J. Trevelyan, Gian-Michele Ratto

**Author notes:** equal contribution (Enrico Pracucci, Robert Graham, Laura Alberio, Gabriele Nardi). equal contribution (Andrew J. Trevelyan, Gian-Michele Ratto).

## Abstract

The main inhibitory synaptic currents, gated by gamma-aminobutyric acid (GABA), are mediated by Cl^-^-conducting channels^1-3^, and are therefore sensitive to changes in the chloride electrochemical gradient. GABAergic activity dictates the neuronal firing range^4,5^ and timing^6-9^, which in turn influences the rhythms of the brain, synaptic plasticity, and flow of information in neuronal networks^7,10-12^. The intracellular chloride concentration [Cl^-^]_i_ ^13^is, therefore, ideally placed to be a regulator of neuronal activity. Chloride levels^13^ have been thought to be stable in adult cortical networks, except when associated with pathological activation^14-17^. Here, we used 2-photon LSSmClopHensor imaging^14^, in anaesthetized young adult mice, to show a large physiological circadian fluctuation of baseline [Cl^-^] inside pyramidal cells, equating to a ∼15mV positive shift in E_Cl_ at times when mice are typically awake (midnight), relative to when they are usually asleep (midday). The change in pyramidal [Cl^-^]_i_ alters the stability of cortical networks, as demonstrated by a greater than 3-fold longer latency to seizures induced by 4-aminopyridine at midday, compared to midnight. Importantly, both [Cl^-^]_i_ and latency to seizure, in night-time experiments, were shifted in line with day-time measures, by inhibition of NKCC1. The redistribution of [Cl^-^]_i_ reflects circadian changes in surface expression and phosphorylation states of the cation-chloride-co-transporters, KCC2 and NKCC1, leading to a greatly reduced chloride-extrusion capacity at night (awake period). Our data show how changes in the biochemical state of neurons may be transduced into altered brain states. (235 words)

---

Fast synaptic inhibition in mammalian brains is subserved primarily by GABA acting upon ionotropic GABA_A_ receptors (GABA_A_-Rs). GABA_A_-Rs are permeable to both chloride and bicarbonate, at a ratio of about 4:1, meaning that the main determinant of the GABA reversal potential (E_GABA_) is the chloride electrochemical gradient^**1-3**^. Intracellular chloride levels are relatively high early in development^14,18,19^, underlying the phenomena of giant depolarizing potentials in prenatal cortex. After birth, the levels drop and adult intracellular [Cl^-^] in neurons is generally considered to lie around 10 mM, setting E_GABA_ substantially below action potential threshold. Chloride levels may rise acutely, following short bursts of GABAergic synaptic activity^17,20^, and can also be loaded into neurons artificially^9,21^, but in both cases, the baseline chloride levels are rapidly re-established through the action of KCC2^9,18^. The potential impact of varying [Cl^-^]_i_ is readily apparent from the suprachiasmatic nucleus (SCN), the one site where significant modulation of neuronal [Cl^-^]_i_ has been demonstrated^13^. SCN neurons receive a strong tonic GABAergic drive, which strongly suppresses neurons during the night, when [Cl^-^]_i_ is low, but leads to strong activation during the day, when [Cl^-^]_i_ is high. This physiological modulation thus underlies the circadian switching of activity in the SCN. Elsewhere in the adult brain, including cortical networks, it was thought that baseline [Cl^-^]_i_ is only raised in association with pathophysiology^9,14-16,22-26^. The recent development of imaging techniques to monitor intracellular [Cl^-^], non-invasively, in living animals^14^, has provided the opportunity to address this preconception about cortical chloride levels.

We introduced the pH and chloride biosensor LSSmClopHensor^14,27^ into neocortical pyramidal cells, in young adult mice. This was achieved in two different ways, in two different laboratories (Pisa and Newcastle). The first labelling strategy (Pisa) was by *in utero* electroporation at gestational day 15.5, using a plasmid encoding LSSmClopHensor under the CAG promoter. This reliably yielded extensive transfection of layer 2/3 pyramidal cells. The second strategy (performed at Newcastle) utilized a viral vector to deliver a floxed construct of LSSmClopHensor, which was injected into the left cerebral ventricle of neonatal P0-P1 mouse pups that express Cre-recombinase under the Emx1 promoter.

Mice were allowed to survive until young adulthood (1–4-month age), being kept on a 12-hour light / 12-hour dark cycle, switching at 7:00 and 19:00. At least 3 days prior to all experimental procedures, we moved the mice into a quiet room, where they were checked just once a day to minimize disturbance to their diurnal cycle. During this time, their activity patterns were video-monitored. In all cases, mice showed a peak of activity in the hour immediately after the lights were turned off, but were typically active throughout the dark period, albeit with short periods of quiescence. In contrast, the mice rarely moved from their nest during the light periods. This diurnal behavioural pattern corresponds well to that of mice in the wild (Extended data figure 1).

We performed *in vivo* LSSmClopHensor imaging of layer 2/3 pyramidal neurons, under anaesthesia, at 4 different times of day: 9:00, 12:00, 19:00 and 23:00 (operations were started within half an hour of these times, and imaging was performed always within 2 hours of the start of the operation). The procedure involved performing a craniotomy over the injected hemisphere, which was covered with a glass coverslip and cemented in place, prior to transferring the animal onto a 2-photon microscope imaging platform. We collected data from various fields of view, between 90-250 µm deep to the pial surface, which corresponds approximately to layers 2-3. In total, we imaged 2575 neurons, from 29 different mice (Figure 1); importantly, the results at the two different centres (which used different transfection strategies for introducing LSSmClopHensor; see above) gave identical results (Extended data figure 2) and the data were therefore pooled. We found a marked variation in [Cl^-^] inside pyramidal cells, over the 24-hour period, with the lowest values occurring around midday (median [Cl^-^]_i_=10.5 mM), in the middle of the mice’s sleep, and the highest values at midnight (median [Cl^-^]_i_=19.0 mM; day vs night, p < 0.005, Mann-Whitney test), when the mice were very active (Figure 1a,b,c). Interestingly, this is the exact opposite cycle to that seen in neurons in the SCN, where [Cl^-^]_i_ is high during the day, and low at night^13^.

**Figure 1.**
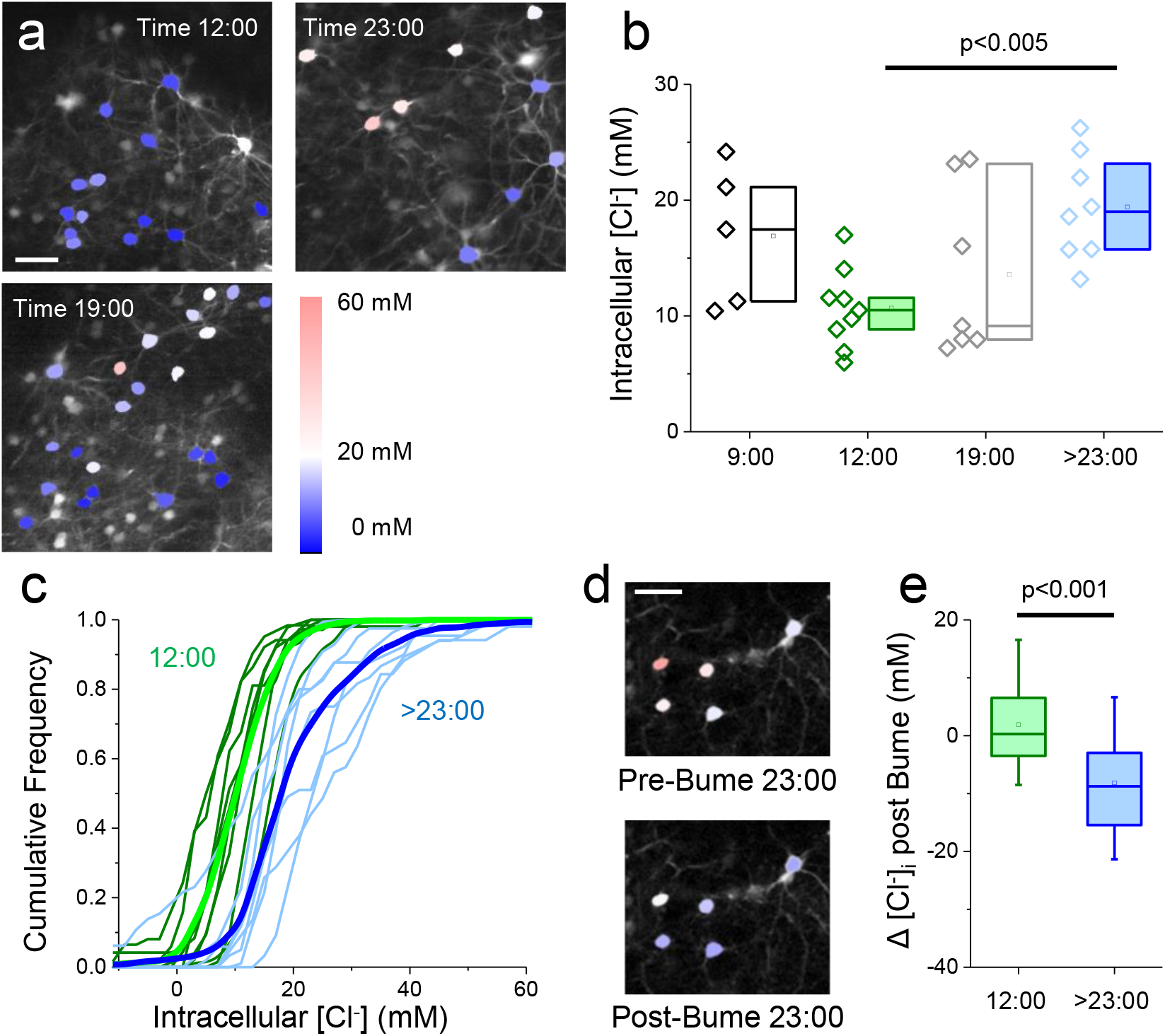
Diurnal change of intracellular Chloride in cortical pyramidal neurons. **a)** Three representative fields of view, showing pyramidal cells expressing LSSmClopHensor, in layers 2/3 of visual cortex, in mice aged ∼1 month. Mice were imaged, under terminal anaesthesia, at the indicated times. Calibration bar 40 μm. **b**) Quantification of [Cl]_i_ in the entire data set. Each symbol is the median value computed from a single animal. The boxplot represents the first and third interquartile, mean and median. Data are obtained from: time 9:00, 5 mice and 159 neurons; time 12:00, 9 mice and 810 neurons; time 19:00, 7 mice and 390 neurons; time >22:30, 8 mice and 1216 neurons. Mann-Whitney test. **c**) Cumulative distribution for each mouse imaged around midday (green traces) and at night (blue traces). The thick lines show the mean distributions of the two groups. **d**) Representative fields imaged at night before and 40 min after superfusion with 50 μM Bumetanide: [Cl]_i_ decreases in all neurons in the field. Calibration bar 40 μm. **e**) Change of [Cl^-^]_i_ measured in the same neuron before and after superfusion of the cortex with Bumetanide. Cumulative data from 4 (188 neurons) and 5 mice (137 neurons) imaged at midday and after 22:30 respectively. The boxplot indicates the first and third interquartile, while the whiskers indicate the 10 and 90 percentiles. Mann-Whitney test.

At dawn and dusk (9:00, 19:00), pyramidal [Cl^-^]_i_ showed intermediate values, but also, interestingly, the largest inter-animal variability, possibly indicating phase differences of the diurnal cycle between the various mice. There was negligible fluctuation in pH across the 4 different imaging times (Extended data figure 3). Importantly, the dispersion of [Cl^-^]_i_ values, within each mouse, was minimal during the day but larger during the night (figure 1c). Notably, in a significant fraction of neurons [Cl^-^]_i_ was larger than 25 mM, which renders E_GABA_ (∼-41mV) more depolarizing than the threshold potential for Na voltage-dependent channels. These data suggest that during the active phase, intracellular [Cl^-^] paints a complex landscape, with a mosaic-like distribution of neurons exhibiting both depolarizing and hyperpolarizing E_Cl_.

Elevated [Cl^-^] within pyramidal neurons, in early postnatal life, can be countered by the focal application of bumetanide, an inhibitor of the Na^+^-K^+^-Cl^-^-cotransporter, NKCC1^14^. We therefore examined whether the nocturnal elevation of [Cl^-^]_i_ in our mice could similarly be altered. Local cortical perfusion of bumetanide had no effect when applied during the day (median change in [Cl^-^]_i_ before and after Bumetanide, ΔCl = 0.29 mM) but drastically reduces [Cl.]_i_ at midnight (median ΔCl = 8.7 mM, Figure 1d,e), suggesting that this variation in [Cl^-^]_i_ through the day reflects a shift in relative activity of NKCC1 and the other cation-chloride cotransporter found in these cells, KCC2.

We next examined what consequences these changes in intracellular [Cl^-^] might have for the cortical network. Raised chloride levels have previously been associated with epileptic pathophysiology^9,14-16,22-26^, so we examined whether there were differences in epileptic susceptibility at midday and midnight, associated with the differences in the pyramidal chloride levels. We made a focal 4-aminopyridine (4-AP) injection (500 nl, 15 mM in saline) directly into the neocortex in urethane-anaesthetized mice, and recorded the evolving neocortical field potential (LFP) in experiments performed at either midnight, or midday (figure 2a,b). Again, the results in both laboratories were in close alignment, and the data were pooled. Mice were recorded for at least 10 min, to ensure a stable electrophysiological baseline, with slow wave oscillations including transient bursts of activity, characteristic of the UP states of non-REM sleep (see details in Extended data figure 4). After 4-AP microinjection, in the midnight experimental group, we observed a rapid build-up of pathological discharges, culminating in a full electrographic seizure in all 6 mice, with a latency of between 7-29 mins after the injection (mean = 17.9 ± 9.5 mins SD, figure 2c and Extended data figure 4). In contrast, identical infusions of 4-AP into neocortex in the midday experimental group induced electrographic seizures in only 3 out of 12 mice (25% of experiments; recordings were terminated after 1 hour after 4-AP injection; Mann-Whitney test vs night untreated, p < 0.005). Assigning the latency for the 75% of recordings that did not progress to seizure, in the midday experiments, as 60 mins (when the recordings were stopped), yields a minimum average of 52 mins, representing at least a 3-fold increase in latency over the midnight experiments. Moreover, if the midnight experimental mice were pretreated with bumetanide, applied directly to the cortical surface for 40 min before 4-AP microinjection, 5 out of 6 recordings (83%) failed to progress to a seizure inside 1hr, and the 6^th^ had a small seizure after 52mins (Mann-Whitney test, night bumetanide vs night untreated, p < 0.005). A different approach, quantifying, instead, the cumulative pathological activity in the three experimental groups (figure 2d,e, and Extended data figure 4, see Methods and Extended data figure 5 for details), confirmed these findings, with a great excess of pathological activity in the midnight recordings, compared to both the midday, and the midnight with bumetanide recordings (Mann-Whitney tests: day vs night, p < 0.02; night vs night/bumetanide, p < 0.01; day vs night/bumetanide, p = 0.59). Thus, blocking NKCC1, using focally applied bumetanide, realigned the cortical excitability at midnight to that shown in the midday experiments, as it did for the intracellular [Cl^-^] levels.

**Figure 2.**
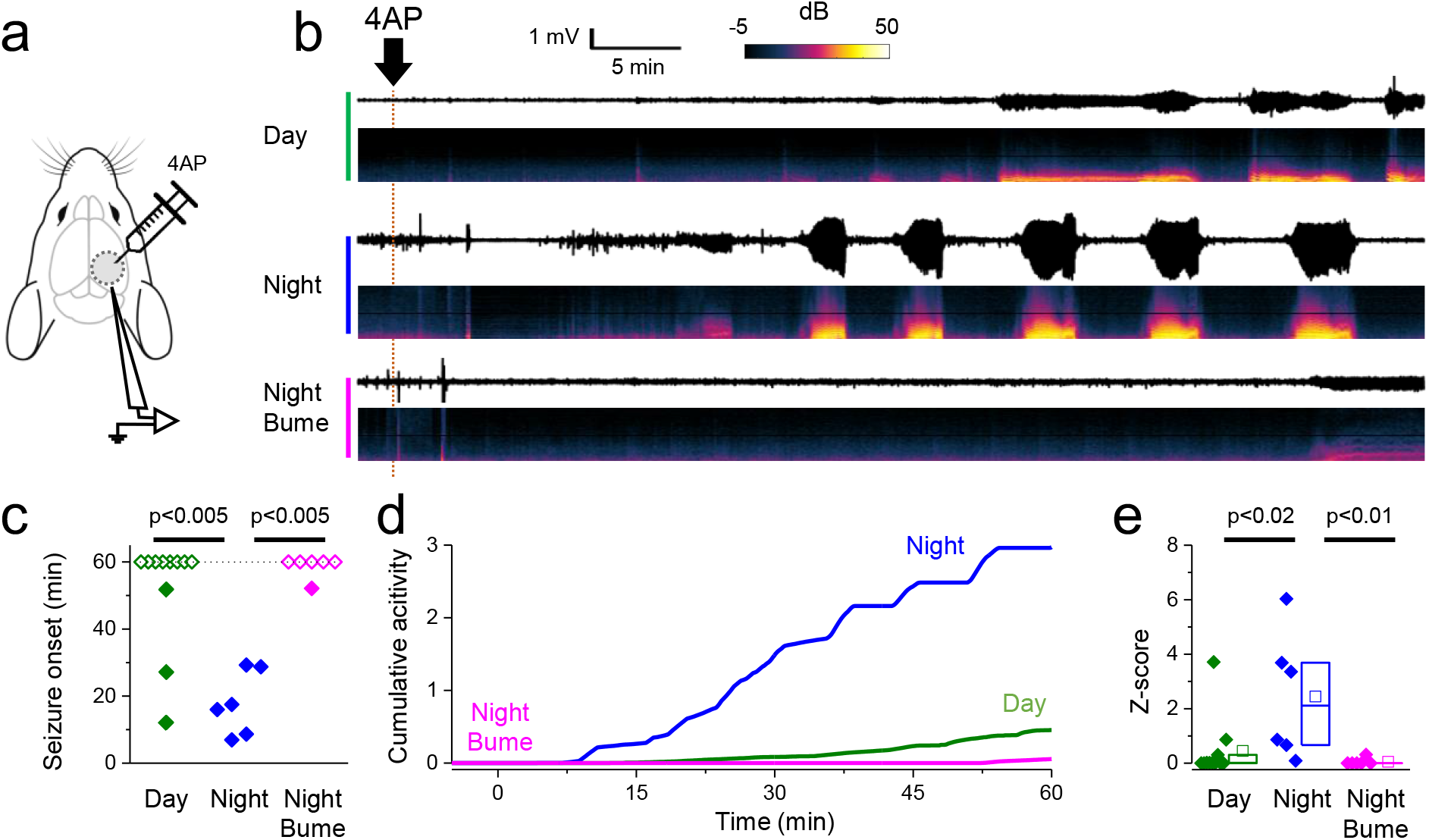
Diurnal change of neuronal excitability related to the Chloride shift. **a)**The LFP is recorded in the occipital cortex under urethane anaesthesia. After attaining a stable baseline, a 0.5 μl injection of 4AP was performed at 1 mm distance from the recording electrode.**b)** Representative traces lasting about 1 hr showing the LFP (black traces) and the relative spectrogram recorded during baseline and after 4AP injection for three mice recorded at midday, midnight and at midnight after a 40 min superfusion with 50 μM Bumetanide. The timing of the beginning of the injection is represented by the orange dotted line. **c**) Lag between the beginning of the 4AP injection and the first recorded epileptiform burst (Wilcoxon rank test, see Methods for the criteria used for burst identification). Empty symbols indicate mice that presented no bursts in a one hour period following the injection. **d**) Mean of the cumulative activity recorded in the three experimental groups. **e**) Z score at 1hr after 4AP microinjection. Box plots represent median and first and third interquartile, Mann-Whitney test.

These physiological data suggest that the diurnal cycle of [Cl^-^]_i,_ and of neuronal excitability, is linked to changes in the relative contribution of NKCC1 and KCC2 to chloride regulation. These are the only chloride-cation cotransporters expressed in neurons and their function is known to be bidirectionally regulated by protein kinases (KCC2 activity is enhanced by phosphorylation of Ser940 ^19,28^ and inhibited by phosphorylation of Thr1007^29^; NKCC1 activity is enhanced by phosphorylation at Thr203 / Thr207 / Thr212^30^). Therefore, we examined the expression levels and phosphorylation state of KCC2 and NKCC1 during the diurnal cycle (figure 3). We first utilized whole-cell biotinylation followed by immunoblotting of neocortical tissue that had been flash-frozen following an acute dissection from freshly sacrificed mice at either midday or midnight. We detected no difference in the total KCC2 protein levels (Mann-Whitney test, n = 4 mice; p = 0.82), but there was a significant reduction in the surface KCC2 protein levels (by 50.2%, Mann-Whitney test, n = 4 mice; p < 0.05) in night mice versus day mice, indicating that there was a marked redistribution of the protein away from the plasma membrane at night. We further examined whether there were alterations of phosphorylation state of either KCC2 or NKCC1 between day and night, indicative of changes in functional activity. We found that KCC2 showed no alteration in phosphorylation at Ser940 (n = 4 mice; p = 0.98), but a significant increased level of KCC2 Thr1007 phosphorylation (by 53.5%, n = 4 mice; p < 0.04), and NKCC1 Thr203/Thr207/Thr212 phosphorylation (by 221.2%, Mann-Whitney test, n = 4 mice; p < 0.05) was detected. These data therefore provide three different mechanisms which would be expected to lead to increased intracellular [Cl^-^] at night, namely (1) a decreased in the cell surface expression of KCC2, (2) phosphorylation of KCC2 at sites which reduces its activity, (3) phosphorylation of NKCC1 at sites which enhances its activity. These biochemical changes thus provide a coherent explanation for the increase in intracellular [Cl^-^] observed during night-time, when the mice are most active.

**Figure 3.**
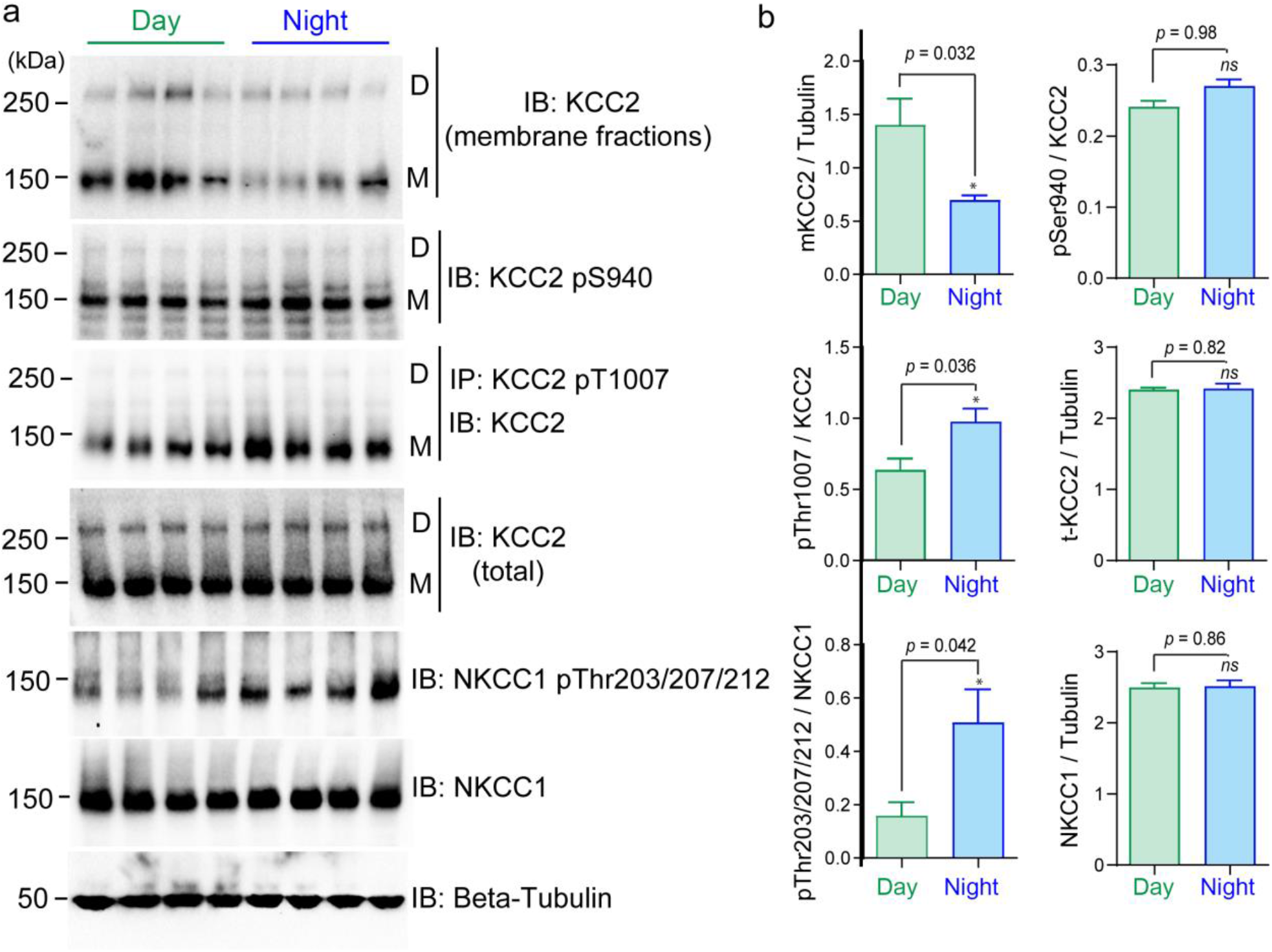
Changes in KCC2 membrane stability and the phosphorylation of KCC2 and NKCC1 in cerebral cortex during the diurnal cycle. **a)** Top panel shows biotinylated surface fraction and total protein expression of KCC2 in membrane fractions prepared from flash-frozen neocortex at different times of day. For following panels: harvested cerebral cortex lysates were subjected to biotinylated surface fraction and total protein expression of KCC2, or immunoprecipitation (IP) with the indicated KCC2 Thr^1007^ phospho-antibody and the immunoprecipitates were then immunoblotted with the indicated specific KCC2 antibody. Cerebral cortex lysates were also subjected to immunoblot analysis with the indicated total and phospho-specific antibodies. Both KCC2 dimers (D) and KCC2 monomers (M) are indicated. Molecular masses are indicated in kDa on the left-hand side of the Western blots. **b**) The right panel shows quantification of the results of the Western blots, as assessed by Mann-Whitney (n=4, error bars represent the mean ± SEM., four independent experiments). The quantification (ratio calculation) is based on (phospho-dimeric KCC2 + phospho-monomeric KCC2) / (total dimeric KCC2 + monomeric KCC2), or Total KCC2 / beta-Tubulin. ***, *p*<0.001; **, *p*<0.01; *, *p*<0.05; ns - not significant.

Collectively, our data show a surprisingly large physiological fluctuation, over the course of a day, in intracellular [Cl^-^] inside cortical pyramidal cells, which is consistent with changes in the phosphorylation state and expression levels of the key chloride-cation cotransporters, suggestive of a causal link. Further evidence for this causal relationship is provided by the effects of bumetanide, which drives an acute change of the midnight pattern towards the midday pattern. The daily flux in pyramidal [Cl^-^]_i_ mirrors that seen in SCN neurons, except that the oscillations at the two sites are out of phase, indicating the potential for cell-class specific modulation with combinatorial consequences for network rhythms. The fluctuations in pyramidal [Cl^-^]_i_ are also associated with large changes in the excitability of the network at different times of the day, consistent with the well-recognized clinical phenomenon of circadian clustering of seizures^31-34^.

Our data add to the growing list of diurnal fluctuations in neuronal function, including changes in firing rates^35-37^, in the number, strength and structure of synapses^38-41^, metabolic state^42^, gene expression and protein phosphorylation^43,44^. Given the direct coupling of the changes in [Cl^-^]_i_ to the biochemical state of neurons, and the well-established role of synaptic inhibition in shaping all manner of neuronal activity^45-51^, we suggest that the changes we describe here, are among the root determinants of the distinct neuronal activity patterns that constitute the cycle of brain states through the day.

## Methods

### Animals and housing conditions

All animal work was performed according to the guidelines of the Home Office UK and animals (Scientific Procedures) Act 1986, and to the protocol 211/2020-PR approved by the Ministero della Salute (Italy). Mice were kept in a 12-hour light / dark cycle, with light being turned on at 7:00 and turned off at 19:00. Food and water were provided ad libitum. Chloride imaging was performed on both CD1 and C57 mice and the electrophysiology was performed on C57 mice only. Most experiments were performed at both sites (Pisa / Newcastle), and data were pooled.

### Production and purification of the AAV8-EF1a-DIO-LSSmClopHensor vector

The virus was produced by the triple transfection method, using 293AAV cells (Cell Biolabs). 293AAV cells were expanded in D-MEM supplemented with 10% FBS, L-glutamine and Pen-Strep, to then transfect 40 × 150 mm plates. For each 150 mm plate, 1.5 × 10^7^ cells were plated the day before the PEI-mediated transfection (Polysciences). For each plate were used 30μg of pAdDeltaF6 helper plasmid (Addgene plasmid # 112867), 15μg of pAAV2-8 Rep-Cap plasmid (Addgene plasmid # 112864), and 15μg of pAAV-EF1a-DIO-ChloPhensor. Sixteen hours after transfection, medium was replaced with D-MEM supplemented with 5% FBS, L-glutamine and Pen-Strep. 72 hours after transfection, cells were scraped and collected by centrifugation.

The virus was purified as described previously^52^. Briefly, the cell pellet was resuspended in sodium citrate pH 8.05 (100mM) using 1.44ml of it + 2.25 of H_2_O per ml of cell pellet. Cells were then sonicated, and 1M magnesium chloride was added to a final concentration of 1.6mM. 200U of benzonase (Sigma-Aldrich) per ml of cell pellet were added and incubated 1h at 37°C. After the benzonase reaction, 3.06ml of citric acid (100mM) + 2.25ml of H_2_O per ml of cell pellet were added to clarify the crude extract by protein flocculation. The extract was then centrifuged at 4500g at RT for 10 minutes. The supernatant was recovered and filtered through a 0.22μm PES filter (Millipore), and directly applied to a HiPrep SP HP 16/10 column (Cytiva) using an AKTA start (Cytiva) connected to a fraction collector Frac30 (Cytiva).

The virus was diafiltrated (1:160000) and concentrated (to final volume of ∼1ml) using Protein Concentrators PES, 100K MWCO (Thermo Scientific, Pierce). The final buffer for the virus was: dPBS + 5% Glycerol + 0.001 Pluronic F-68 (Invitrogen). The AAV titre (1.71 × 10^13^ GC/ml) was determined by qPCR using a standard curve of linearised pAAV-EF1a-DIO-LSSmChloPhensor plasmid as reference.

### LSSmClopHensor transfection

LSSmClopHensor was transduced by *in utero* electroporation in CD-1 mice at embryonic day 15.5 with a custom-made triple electrode to transfect neuronal progenitors of layer 2/3 pyramidal neurons of the visual cortex. Details have been presented elsewhere^53,54^ Transfection using the AAV-LSSmClopHensor viral vectors was performed on neonatal mice pups (postnatal day 0-1) expressing Cre-recombinase under the Emx1 promoter, following a previously described protocol^9^. In short, pups were anaesthetized using isoflurane (2% in O_2_ delivered at 0.4 L/min). The scalp and skull were perforated using a 23G needle, approximately midway between bregma and lambda, and roughly 1 mm lateral to the midline, on the left side. A Hamilton nanofill syringe needle (36G, World Precision Instruments) was then guided through the hole, to a depth of 1.7mm, aiming for the lateral ventricles, and 500nl of AAV8 EF1a DIO LSSmClopHensor viral vector was delivered at that depth, and further boluses also delivered, at 2 min intervals, also at 1.4, 1.1 and 0.8 mm depth. The needle was left in situ for at least 3 mins after the final bolus injection. Mice were allowed to recover fully, while being kept warm, before being returned to the mother.

### LSSmClopHensor imaging

All imaging experiments were performed as acute terminal experiments on young mice (age 1-4 months), which had been transfected previously with LSSmClopHensor. Preparatory surgery followed two slightly different protocols in the two centres. In Pisa, mice were anesthetized with 2,2,2-tribromoethanol (Avertin) (i.p. 0.02 ml/g body weight) whereas in Newcastle, urethane anaesthesia was used (i.p. 1.6 g/kg), and supplemented with low level of isoflurane inhalation (5% in 0_2_ for induction, and then lowered to <1% in O_2_ at 0.6-1 L/min, for maintenance during the craniotomy, but discontinued for more than 10minutes, prior to imaging). In both cases, a 3 mm craniotomy was performed over the occipital cortex and covered with a 3mm-coverslip. In some experiments, the coverslip was left partially open on a side to allow for the bumetanide treatment. Imaging was performed immediately after surgery.

All imaging sessions were performed in identical fashion, at four different times during the day, around 7:00 / 12:00 / 19:00 / >23:00. Once mice were fully anaesthetized, they were positioned underneath a 2-photon microscope. The injections sites were identified from the vasculature landmarks, and labelling confirmed quickly, while identifying optimal fields of view. Images were acquired at 5 different excitation wavelengths (800 / 830 / 860 / 910 / 960 nm) and collected through green and a red emission filters (527/70 nm and 607/70 nm BrightLine®). Post hoc analysis of the images was performed using a graphical user interface (GUI), implemented using MATLAB software (Mathworks), that automated finding cell somata, to collect red/green fluorescence ratios at the 5 different wavelengths, and compute intracellular pH and [Cl^-^] values for each cell, based upon earlier calibration experiments performed using ionophore-permeabilized HEK cell and neuronal cultures^14^.

The two centres used two different imaging setups. Pisa: Bruker Ultima with a Coherent Ultra II laser and an Olympus 20X, NA 1.05 objective (water immersion). Newcastle: Bergamo II 2-photon microscope, with a MaiTai laser (SpectraPhysics) and a Nikon 16X, NA 0.8 (water immersion). Of note, both microscopes were equipped with identical filters on the emission pathway.

### Seizure susceptibility

These experiments were duplicated in both Pisa and Newcastle, following the same broad protocol, albeit with minor differences as indicated below. *In vivo* electrophysiological experiments were performed on acutely prepared C57BL/6 mice, under urethane anaesthesia (i.p. 1.6 g/kg). In Newcastle, anaesthesia was supplemented by 0.5%, isoflurane in oxygen (0.4-1 l/min) while the craniotomy was being performed but was discontinued prior to seizure induction (as per the LSSmClophensor imaging experiments), because it has anti-epileptic effects. The mouse was placed in a stereotaxic head-holder and a small craniotomy was performed to reveal the cortex. Recording was performed either using 2 MOhm glass microelectrodes (Pisa), or a multi-electrode array (MEA, Neuronexus – A4×4-4mm-200-200-1250, in Newcastle). In both cases, electrodes were introduced into the cortex to a depth of 0.3 mm, and approximately positioned within the primary somatosensory cortex^55^. The cortex was kept moist with warmed saline throughout recording sessions. LFP signals were amplified and digitised at 10 kHz (Pisa, NPI EXT-02F amplifier acquired with the Neuronexus SmartBox Pro; Newcastle: Plexon AC amplifier and stored using the SortClient software).

Epileptic activity was induced acutely by a single bolus injection of 500 nl of 15 mM 4-aminopyridine in saline (total amount delivered 7.5 nmol), injected at 1 µl/min via a NanoFil needle (36G, World Precision Instruments; 30 s injection, Newcastle) or at 0.2 µl/min via a 40 μm glass pipette (Legato 130, KD Scientific microinjector; 2.5 min injection, Pisa), at a distance of 1mm caudal to the recording site (approximately into the anterior end of primary visual cortex).

### Electrophysiology analysis

Extracellular field recordings were analysed to detect pathological discharges using a custom-written code in Matlab2020b (Mathworks, MA, USA) which can be provided upon request. In brief, this used a frequency-domain analysis to detect periods when the LFP power exceeded baseline activity by a designated threshold (Extended data figure 5). The spectrogram (bin width 10 s, with 1 s time shift) of the baseline (the period preceding 4AP injection) was computed and we obtained mean and standard deviation of the spectral power for frequencies between 8-100Hz. The spectrogram of the entire recording was normalized to the baseline mean and standard deviation for each frequency bandwidth individually. To avoid contamination with mains noise, we omitted frequencies between 49-51Hz. This process generated a Z-score spectrogram (Z = (x-mean)/s.d.), in which teach element of the Z-score matrix represented the excess power at a particular frequency, for each time bin. Using a threshold of 4 standard deviations above the baseline mean, we identified all time bins for which any frequency exceeded this threshold. The power integrated in each time bin was summed cumulatively, to estimate the rate and acceleration of pathological activity within the brain slice (Extended data figure 5). Continuous epochs of suprathreshold bins were designated to be single “events”. Seizures were identified as continual large amplitude rhythmic discharges lasting more than 15 s. In fact, most seizures lasted for many tens of seconds, before transitioning into a period of postictal suppression.

### KCC2/NKCC1 analysis

#### Animals

Mice were killed by cervical dislocation, and the brains removed into ice-cold saline, and a rapid dissection was made to isolate right and left neocortical and hippocampal brain tissue samples, which were immediately flash-frozen in liquid nitrogen, and stored subsequently at - 80°.

#### Antibodies

Primary antibodies used were: anti-KCC2 phospho-Ser940 (Thermo Fisher Scientific, cat #PA5-95678), anti-(neuronal)-β-Tubulin III (Sigma-Aldrich, cat #T8578), anti-KCC2 total (University of Dundee, S700C), anti-KCC2 phospho-Thr1007 (University of Dundee, S961C), anti-NKCC1 total antibody (University of Dundee, S022D), anti-NKCC1 & NKCC1 phospho-Thr203/Thr207/Thr212 antibody (University of Dundee, S763B). Horseradish peroxidase-coupled (HRP) secondary antibodies used for immunoblotting were from Pierce. IgG used in control immunoprecipitation experiments was affinity-purified from pre-immune serum using Protein G-Sepharose.

#### Buffer for Western Blots

Buffer A contained 50 mM Tris/HCl, pH7.5 and 0.1mM EGTA. Lysis buffer was 50 mM Tris/HCl, pH 7.5, 1 mM EGTA, 1 mM EDTA, 50 mM sodium fluoride, 5 mM sodium pyrophosphate, 1 mM sodium orthovanadate, 1% (w/v) NP40, 0.27 M sucrose, 0.1% (v/v) 2-mercaptoethanol, and protease inhibitors (complete protease inhibitor cocktail tablets, Roche, 1 tablet per 50 mL). TBS-Tween buffer (TTBS) was Tris/HCl, pH 7.5, 0.15 M NaCl and 0.2% (v/v) Tween-20. SDS sample buffer was 1X NuPAGE™ LDS Sample Buffer (NP0007, Invitrogen™), containing 1% (v/v) 2-mercaptoethanol. Protein concentrations were determined following centrifugation of the lysate at 16,000 x g at 4 °C for 20 minutes using the using a Pierce™ Coomassie (Bradford) Protein Assay Kit (23200, Thermo Scientific) with bovine serum albumin as the standard.

#### Surface biotinylation in cerebral cortex

Biotinylation studies were performed as previously described^56^ with modifications. Mice were killed by cervical dislocation and the brain rapidly extracted and placed in a cold (∼4°C), oxygenated sucrose-based solution, comprising (in mM): sucrose (189), d-glucose (10), NaHCO_3_ (26), KCl (3), MgSO_4_ (5), CaCl_2_ (0.1) and NaH_2_PO_4_ (1.25). Horizontal sections (400 µm) were made from 5–7-week-old wild-type animals (C57Bl/6j, Janvier) using a vibrating blade microtome (VT1200, Leica Microsystems, Wetzlar, Germany) in NMDG based cutting solution (in mm): NMDG (93), HCl (93), KCl (2.5), NaH_2_PO_4_ (1.2), NaHCO_3_ (30), HEPES (20), glucose (25) ascorbic acid (5), sodium pyruvate (3), MgCl_2_ (10), CaCl_2_ (0.5), saturated with 95% O_2_/5% CO_2_, pH 7.4, 300 mOsm). After cutting, the slices were immediately removed to a holding chamber containing oxygenated (95% O_2_–5% CO2) artificial cerebrospinal fluid (aCSF) comprising (in mm): NaCl (124), KCl (3), NaHCO_3_ (24), MgSO_4_ (1), d-glucose (10) and CaCl_2_ (1.2). The slices were gradually warmed to ∼37°C (for 30 min) and then maintained at room temperature (∼20°C, for at least another 30 min) until ready for use. The slices were transferred into bubbled ice-cold aCSF containing 1 mg/ml EZ-Link™ Sulfo-NHS-Biotin (21217, Thermo Scientific) with gentle rotation for 45 min at 4 °C. Excess biotin was quenched using 1 M glycine in ice-cold aCSF for 10 min, and then the slices were rinsed once in ice-cold aCSF and snap frozen on dry ice. The cerebral cortex was micro-dissected and immediately lysed and homogenized in lysis buffer. The samples were centrifuged at 16,000 x g at 4 °C for 20 min, the supernatant was collected, and protein content was determined using a Pierce™ Coomassie (Bradford) Protein Assay Kit (23200, Thermo Scientific). 40 µg of protein was loaded onto 100 µl of 50% slurry of Pierce™ NeutrAvidin™ UltraLink™ Resin (53150, Thermo Scientific), made up to a total volume of 400 µl in lysis buffer and rotated for 2hr at 4 °C. The beads were recuperated by centrifugation and thoroughly washed four times in lysis buffer, and after the last wash the beads were incubated in ×2 SDS sample buffer containing 10% β-mercaptoethanol at 37 °C for 1hr. The protein samples were run on pre-cast Bis-Tris gels (NuPageTM 4–12% gradient gels, NP0322, Invitrogen™) and immunoblotting was performed. Analysis was performed using ImageJ by normalizing the amount of surface KCC2 to the amount of β-tubulin in the non-biotinylated fraction.

#### Immunoblot and phospho-antibody immunoprecipitation analyses

Cerebral cortex lysates were subjected to immunoblot and immunoprecipitation as previously described^57^. Cerebral cortex lysates (20 µg) were boiled at 75 °C in sample buffer for 10 minutes, resolved by 7.5 % sodium dodecyl sulfate polyacrylamide-gel electrophoresis and electrotransferred onto a polyvinylidene difluoride membrane. Membranes were incubated for 30 min with TBST (Tris-buffered saline, 0.05% Tween-20) containing 5% (w/v) skim milk. Blots were then washed three times with TBST and incubated for 1hr at room temperature with secondary HRP-conjugated antibodies diluted 5000-fold in 5% (w/v) skim milk in TBST. After repeating the washing steps, signals were detected with enhanced chemiluminescence reagent. Immunoblots were developed using ChemiDoc™ Imaging Systems (Bio-Rad, Feldkirchen). Figures were generated using Photoshop/Illustrator (Adobe). The relative intensities of immunoblot bands were determined by densitometry with ImageJ software. Calculation of intensity ratios^58^ is based on (phospho-dimeric KCC2 + phospho-monomeric KCC2) /(total dimeric KCC2 + total monomeric KCC2), (total dimeric KCC2 + total monomeric KCC2)/β-tubulin, NKCC1 pThr203,207,212 /NKCC1, NKCC1/β-tubulin, as reported previously^57^. Data were expressed as means SEM. Statistical significance was determined by Wilcoxon-Mann-Whitney sum test and adjusted by Bonferroni correction (GraphPad Prism 7.0, San Diego, CA, USA).

KCCs phosphorylated at the KCC2 Thr^1007^ equivalent residue were immunoprecipitated from clarified clarified cerebral cortex lysates (centrifuged at 16,000 x g at 4 °C for 20 minutes) using phosphorylation site-specific antibody coupled to protein G–Sepharose as previously described^57^. The phosphorylation site-specific antibody was coupled with protein-G–Sepharose at a ratio of 1 mg of antibody per 1 mL of beads in the presence of 20 µg/mL of lysate to which the corresponding non-phosphorylated peptide had been added. Two mg of clarified cell lysate were incubated with 15 µg of antibody conjugated to 15 µL of protein-G–Sepharose for 2 hours at 4°C with gentle agitation. Beads were washed three times with 1 mL of lysis buffer containing 0.15 M NaCl and twice with 1 mL of buffer A. Bound proteins were eluted with 1X LDS sample buffer.

## Extended data

**Extended data figure 1.**
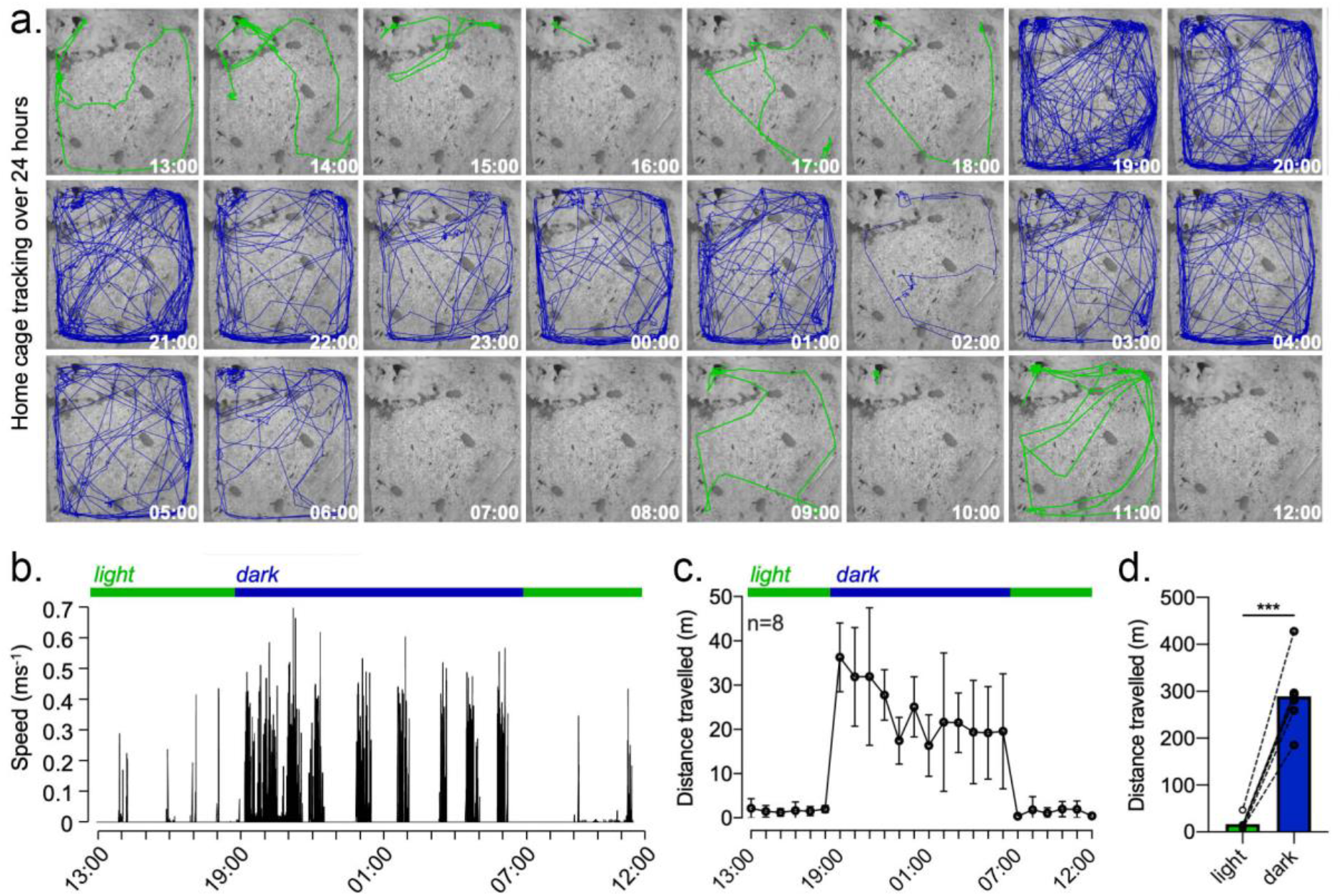
Mice are active almost exclusively during periods of darkness. **a)** Each panel shows the movement of a single mouse during a single hour, as specified, sampled every 2s. Green lines were during the light periods (7:00-19:00), and blue lines during the dark periods (19:00-7:00). **b**) A higher resolution depiction of the movement speed for a single mouse. Note the sudden transformation of activity, almost as soon as the lights are turned off, and while there are subsequently periods of inactivity during nighttime, the activity remains far higher than during the day, when mice are almost entirely inactive. **c, d**) The pooled data show that this pattern was repeated in all mice and is in line with the known behaviour patterns of mice in the wild.

**Extended data figure 2.**
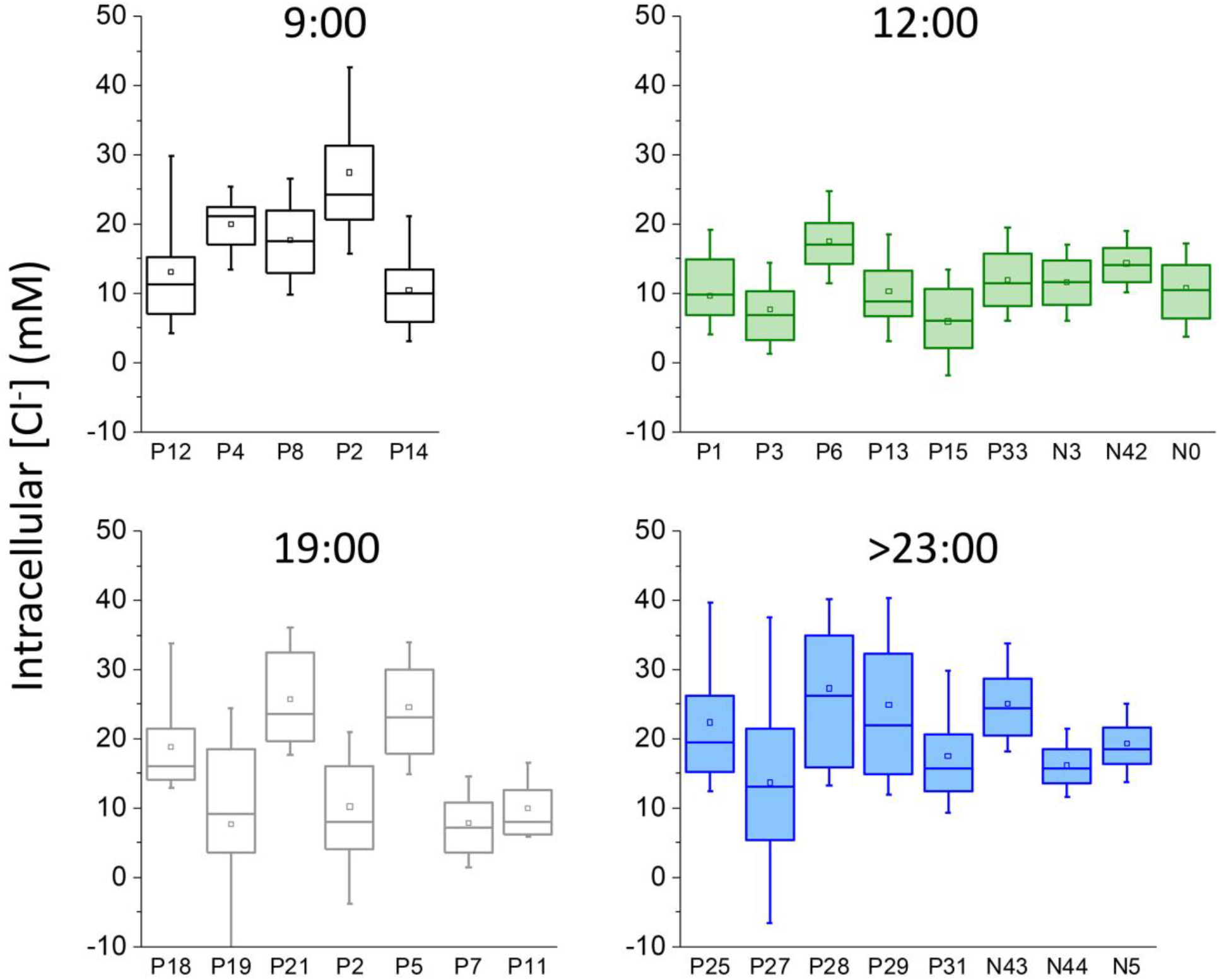
Boxplots of chloride measurements in all animals. The suffix ‘P’ denotes mice imaged in Pisa, and ‘N’ those imaged in Newcastle. Note the consistency between the data sets acquired at the two sites.

**Extended data figure 3.**
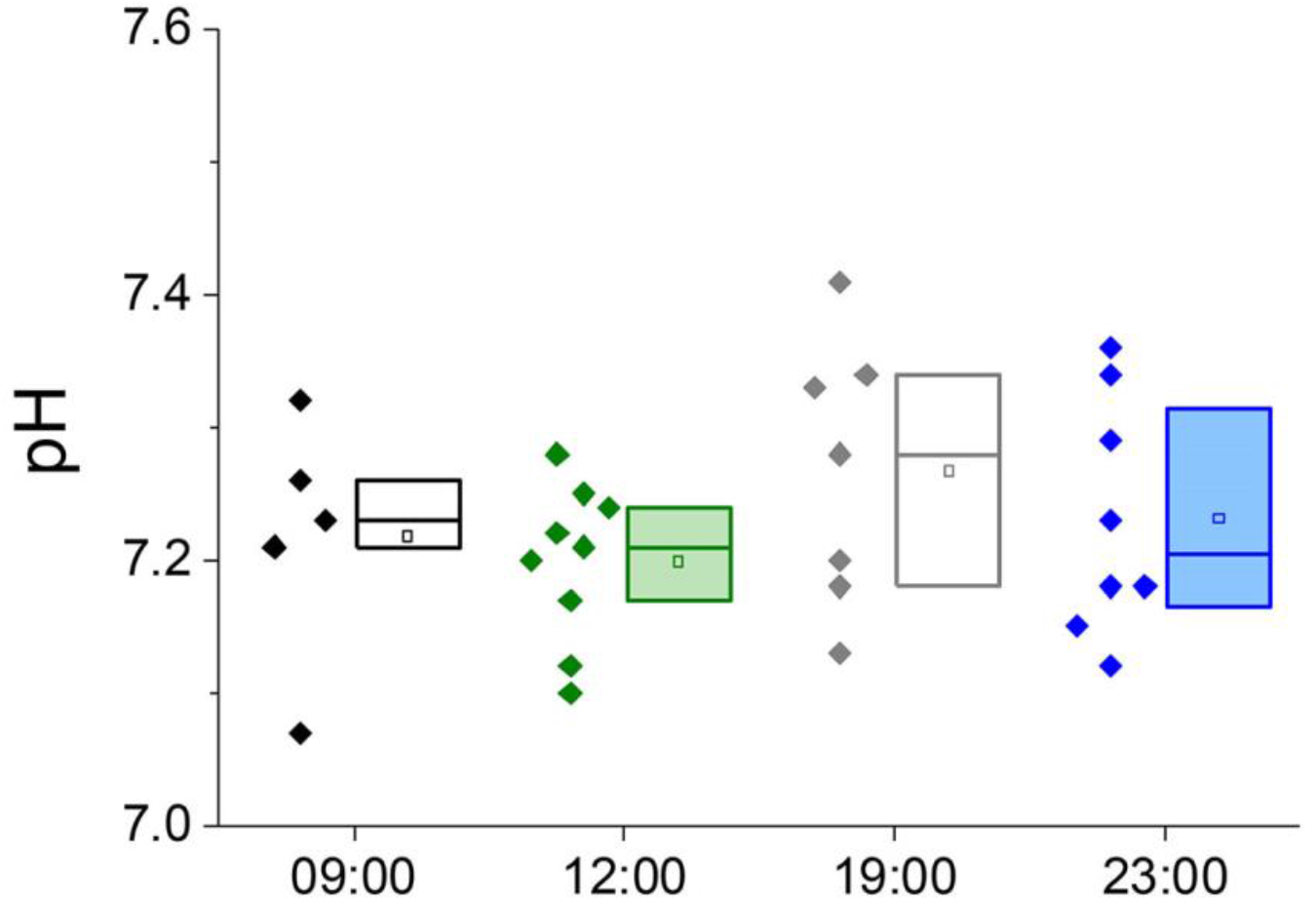
Intracellular pH, imaged using LSSmClopHensor, shows negligible changes over the course of the day. Each data point represents the mean of all pyramidal cells imaged from a single animal, and the box plots represent the intraquartile range, median (line) and mean (open point). Groups are not significantly different (p>0.20, Mann-Whitney test).

**Extended data figure 4.**
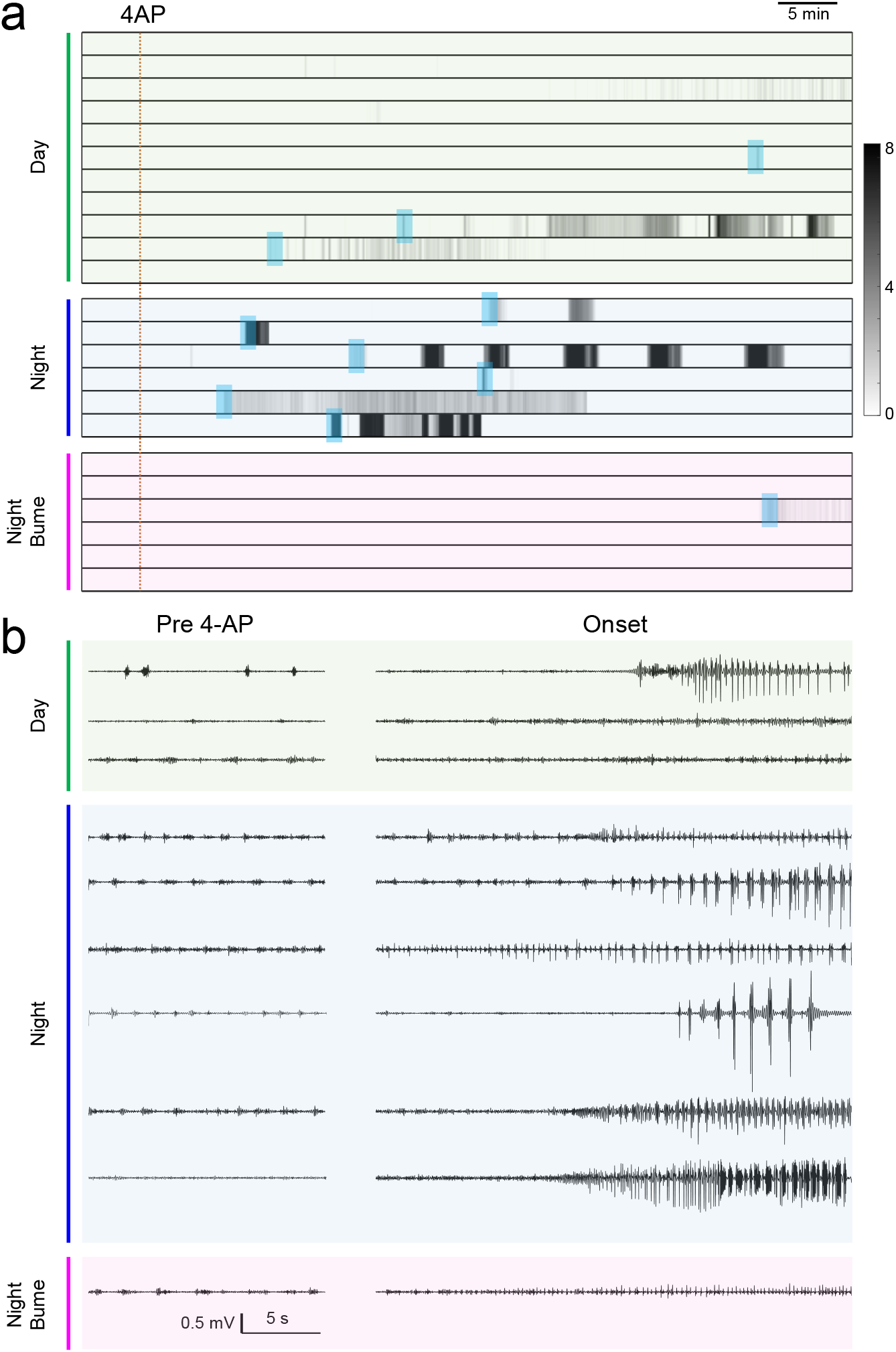
Extended data figure 4 Analysis of the electrophysiology data set. **a)** Raster plots showing the temporal evolution of the Z-score (gray-scale) for all mice included in the study within the three experimental groups: (1) mice recorded during the day, green; (2) at night, blue; and (3) at night after treatment with bumetanide, mauve. The shadowed areas are around the onset of epileptic activity, and these events are enlarged in panel b. **b**) Baseline activity (left) and onset of epileptic activity (right traces). Data have been band passed in the 8-80 Hz band.

**Extended data figure 5.**
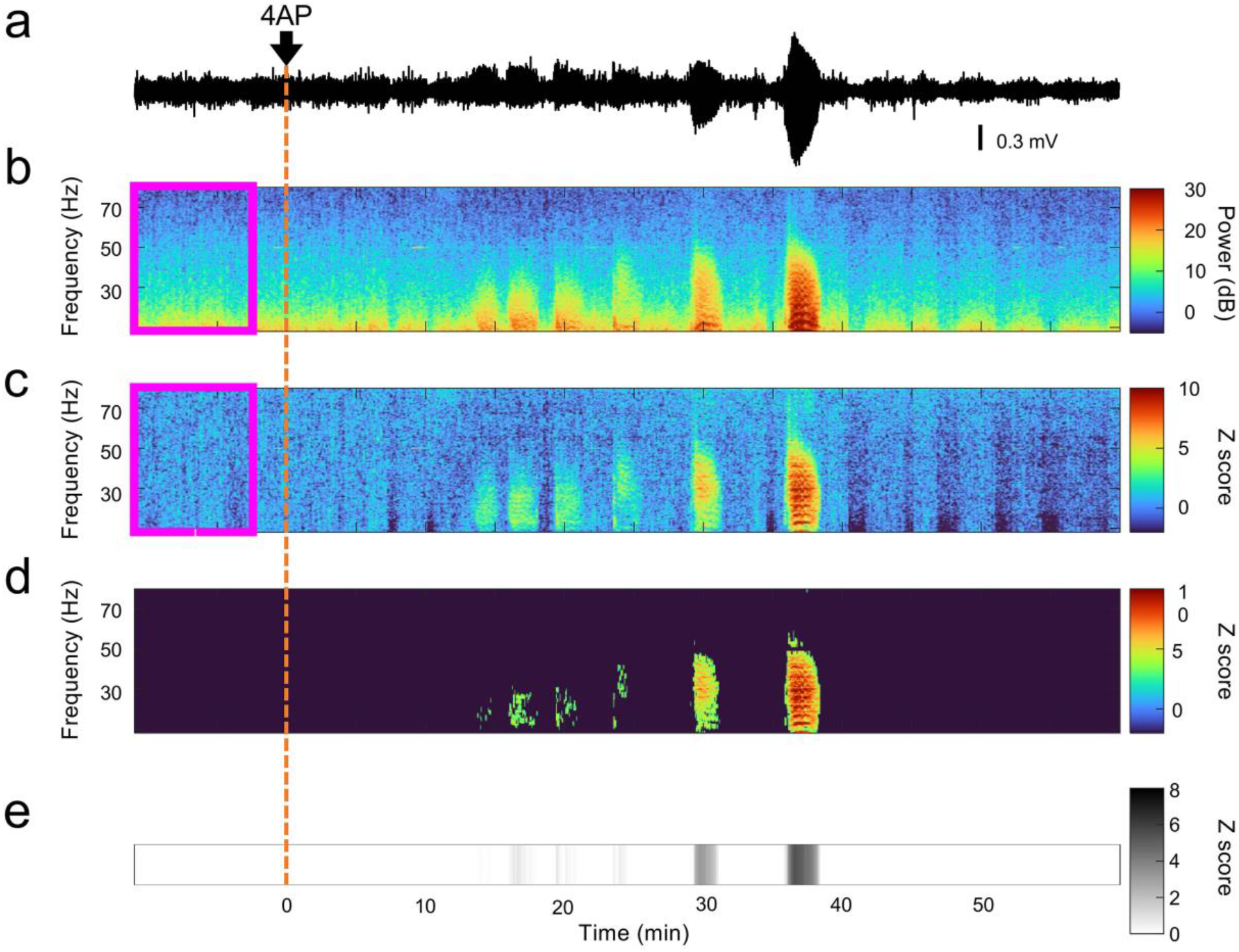
Computation of the Z-scoring of epileptiform activity. **a)** An example trace of the LFP recording, during a midnight 4-AP experiment. 4AP was microinjected over a period of 2 min, starting at time 0. **b**) spectrogram of the same trace, computed using the Matlab function mtspecgramc (time windows of 10 s with steps of 1 s, 5 tapers and time-bandwidth product = 3). Frequencies between 8 Hz and 100 Hz are used. Note that power attenuates at high frequencies. The magenta frame highlights a baseline period before 4AP injection. **c**) Z-transform of the spectrogram in b computed as follows: for each frequency f, we computed the mean (μ_f_) and the standard deviation (σ_f_) of the power during the baseline. Then, for every element of the spectrogram x_ft_, the z transform is obtained with (x_ft_ – μ_f_) / σ_f_. Note that this normalization overcomes the attenuation of the power at high frequencies. **d)** a binary mask is applied to the z-transform of the spectrogram to keep only values higher than 4 (4σ higher than the baseline power). **e**) the z-score time course is computed, summing the z-transform of the spectrogram across all frequencies (columns).

## Contributions and acknowledgements

Study design: GMR, AT. Experiment supervision (Pisa): GMR, SL; (Newcastle): AT. In vivo imaging (Pisa): OC, SL; (Newcastle): RG, LA. Electrophysiology (Pisa): EP, GN, VP, GMR; (Newcastle): RG, LA. Design and production of viral vector: LS, LA. Biochemistry: JZ. Computer code: EP, GN, GMR, DW, RG, AT. Data analysis (imaging): OC, SL, RG, LA. Data analysis (electrophysiology): EP, GN, GMR, RG, DW, AT. Manuscript preparation: AT, GMR with contributions from all authors. We are grateful to Francesca Biondi for animal care. The study was funded by Fondazione Telethon grant GGP19281, Regione Toscana (projects DEMAGING and DECODEE), MIUR (PRIN 20175C22WM), The Royal Society (IEC\NSFC\201094), MRC (MR/R005427/1), BBSRC (BB/P019854/1), and a Newcastle University PhD studentship (RG) and Newcastle University Research Fellowship (LA).

## References

1. Kaila, K. & Voipio, J. Postsynaptic fall in intracellular pH induced by GABA-activated bicarbonate conductance. Nature 330, 163–165 (1987).

2. Bormann, J., Hamill, O.P. & Sakmann, B. Mechanism of anion permeation through channels gated by glycine and gamma-aminobutyric acid in mouse cultured spinal neurones. J Physiol 385, 243–286 (1987).

3. Coombs, J.S., Eccles, J.C. & Fatt, P. The specific ionic conductances and the ionic movements across the motoneuronal membrane that produce the inhibitory post-synaptic potential. J Physiol 130, 326–374 (1955).

4. Pouille, F., Watkinson, O., Scanziani, M. & Trevelyan, A.J. Differential control of pyramidal neurons’ dynamic range by somatic and dendritic inhibition. (Society for Neuroscience, 2008., Washington, DC, 2008).

5. Pouille, F., Watkinson, O., Scanziani, M. & Trevelyan, A.J. The contribution of synaptic location to inhibitory gain control in pyramidal cells. Physiological reports 1, e00067 (2013).

6. Pouille, F. & Scanziani, M. Enforcement of temporal fidelity in pyramidal cells by somatic feed-forward inhibition. Science 293, 1159–1163 (2001).

7. Cobb, S.R., Buhl, E.H., Halasy, K., Paulsen, O. & Somogyi, P. Synchronization of neuronal activity in hippocampus by individual GABAergic interneurons. Nature 378, 75–78 (1995).

8. Buzsáki, G. Rhythms of the brain, (Oxford University Press, Oxford, 2006).

9. Alfonsa, H., et al. The contribution of raised intraneuronal chloride to epileptic network activity. J Neurosci 35, 7715–7726 (2015).

10. Atallah, B.V., Bruns, W., Carandini, M. & Scanziani, M. Parvalbumin-expressing interneurons linearly transform cortical responses to visual stimuli. Neuron 73, 159–170 (2012).

11. Melzer, S., et al. Long-range-projecting GABAergic neurons modulate inhibition in hippocampus and entorhinal cortex. Science 335, 1506–1510 (2012).

12. Buzsaki, G. & Wang, X.J. Mechanisms of gamma oscillations. Annu Rev Neurosci 35, 203–225 (2012).

13. Wagner, S., Castel, M., Gainer, H. & Yarom, Y. GABA in the mammalian suprachiasmatic nucleus and its role in diurnal rhythmicity. Nature 387, 598–603 (1997).

14. Sulis Sato, S., et al. Simultaneous two-photon imaging of intracellular chloride concentration and pH in mouse pyramidal neurons in vivo. Proc Natl Acad Sci U S A 114, E8770–E8779 (2017).

15. Ellender, T.J., Raimondo, J.V., Irkle, A., Lamsa, K.P. & Akerman, C.J. Excitatory effects of parvalbumin-expressing interneurons maintain hippocampal epileptiform activity via synchronous afterdischarges. J Neurosci 34, 15208–15222 (2014).

16. Pavlov, I., Kaila, K., Kullmann, D.M. & Miles, R. Cortical inhibition, pH and cell excitability in epilepsy: what are optimal targets for antiepileptic interventions? J Physiol 591, 765–774 (2013).

17. Staley, K.J., Soldo, B.L. & Proctor, W.R. Ionic mechanisms of neuronal excitation by inhibitory GABAA receptors. Science 269, 977–981 (1995).

18. Rivera, C., et al. The K+/Cl-co-transporter KCC2 renders GABA hyperpolarizing during neuronal maturation. Nature 397, 251–255 (1999).

19. Leonzino, M., et al. The Timing of the Excitatory-to-Inhibitory GABA Switch Is Regulated by the Oxytocin Receptor via KCC2. Cell Rep 15, 96–103 (2016).

20. Thompson, S.M. & Gahwiler, B.H. Activity-dependent disinhibition. I. Repetitive stimulation reduces IPSP driving force and conductance in the hippocampus in vitro. J Neurophysiol 61, 501–511 (1989).

21. Raimondo, J.V., Kay, L., Ellender, T.J. & Akerman, C.J. Optogenetic silencing strategies differ in their effects on inhibitory synaptic transmission. Nat Neurosci 15, 1102–1104 (2012).

22. Huberfeld, G., et al. Perturbed chloride homeostasis and GABAergic signaling in human temporal lobe epilepsy. J Neurosci 27, 9866–9873 (2007).

23. Dzhala, V.I., et al. Progressive NKCC1-dependent neuronal chloride accumulation during neonatal seizures. J Neurosci 30, 11745–11761 (2010).

24. Avoli, M. & de Curtis, M. GABAergic synchronization in the limbic system and its role in the generation of epileptiform activity. Prog Neurobiol 95, 104–132 (2011).

25. Chang, M., et al. Brief activation of GABAergic interneurons initiates the transition to ictal events through post-inhibitory rebound excitation. Neurobiol Dis 109, 102–116 (2018).

26. Cossart, R., Bernard, C. & Ben-Ari, Y. Multiple facets of GABAergic neurons and synapses: multiple fates of GABA signalling in epilepsies. Trends Neurosci 28, 108–115 (2005).

27. Arosio, D., et al. Simultaneous intracellular chloride and pH measurements using a GFP-based sensor. Nat Methods 7, 516–518 (2010).

28. Lee, H.H., et al. Direct protein kinase C-dependent phosphorylation regulates the cell surface stability and activity of the potassium chloride cotransporter KCC2. J Biol Chem 282, 29777–29784 (2007).

29. Rinehart, J., et al. Sites of regulated phosphorylation that control K-Cl cotransporter activity. Cell 138, 525–536 (2009).

30. Zhang, J., et al. Functional kinomics establishes a critical node of volume-sensitive cation-Cl(-) cotransporter regulation in the mammalian brain. Scientific reports 6, 35986 (2016).

31. Karoly, P.J., et al. The circadian profile of epilepsy improves seizure forecasting. Brain 140, 2169–2182 (2017).

32. Karoly, P.J., et al. Circadian and circaseptan rhythms in human epilepsy: a retrospective cohort study. Lancet Neurol 17, 977–985 (2018).

33. Baud, M.O., et al. Multi-day rhythms modulate seizure risk in epilepsy. Nature communications 9, 88 (2018).

34. Karoly, P.J., et al. Cycles in epilepsy. Nat Rev Neurol (2021).

35. Torrado Pacheco, A., Bottorff, J., Gao, Y. & Turrigiano, G.G. Sleep Promotes Downward Firing Rate Homeostasis. Neuron 109, 530–544 e536 (2021).

36. Watson, B.O., Levenstein, D., Greene, J.P., Gelinas, J.N. & Buzsaki, G. Network Homeostasis and State Dynamics of Neocortical Sleep. Neuron 90, 839–852 (2016).

37. Vyazovskiy, V.V., et al. Cortical firing and sleep homeostasis. Neuron 63, 865–878 (2009).

38. Tononi, G. & Cirelli, C. Sleep function and synaptic homeostasis. Sleep Med Rev 10, 49–62 (2006).

39. Liu, Z.W., Faraguna, U., Cirelli, C., Tononi, G. & Gao, X.B. Direct evidence for wake-related increases and sleep-related decreases in synaptic strength in rodent cortex. J Neurosci 30, 8671–8675 (2010).

40. Vyazovskiy, V.V., Cirelli, C., Pfister-Genskow, M., Faraguna, U. & Tononi, G. Molecular and electrophysiological evidence for net synaptic potentiation in wake and depression in sleep. Nat Neurosci 11, 200–208 (2008).

41. Zhou, Y., et al. REM sleep promotes experience-dependent dendritic spine elimination in the mouse cortex. Nature communications 11, 4819 (2020).

42. Natsubori, A., et al. Intracellular ATP levels in mouse cortical excitatory neurons varies with sleep-wake states. Commun Biol 3, 491 (2020).

43. Wang, Z., et al. Quantitative phosphoproteomic analysis of the molecular substrates of sleep need. Nature 558, 435–439 (2018).

44. Bruning, F., et al. Sleep-wake cycles drive daily dynamics of synaptic phosphorylation. Science 366(2019).

45. Prince, D.A. & Wilder, B.J. Control mechanisms in cortical epileptogenic foci. “Surround” inhibition. Arch Neurol 16, 194–202 (1967).

46. Wilson, H.R. & Cowan, J.D. Excitatory and inhibitory interactions in localized populations of model neurons. Biophys J 12, 1–24 (1972).

47. Dudok, B., et al. Alternating sources of perisomatic inhibition during behavior. Neuron 109, 997–1012 e1019 (2021).

48. Pelkey, K.A., et al. Hippocampal GABAergic Inhibitory Interneurons. Physiol Rev 97, 1619–1747 (2017).

49. Lovett-Barron, M., et al. Regulation of neuronal input transformations by tunable dendritic inhibition. Nat Neurosci 15, 423–430 (2012).

50. Isaacson, J.S. & Scanziani, M. How inhibition shapes cortical activity. Neuron 72, 231–243 (2011).

51. Trevelyan, A.J. Do Cortical Circuits Need Protecting from Themselves? Trends Neurosci 39, 502–511 (2016).

52. Potter, M., et al. A simplified purification protocol for recombinant adeno-associated virus vectors. Mol Ther Methods Clin Dev 1, 14034 (2014).

53. Szczurkowska, J., et al. Targeted in vivo genetic manipulation of the mouse or rat brain by in utero electroporation with a triple-electrode probe. Nat Protoc 11, 399–412 (2016).

54. dal Maschio, M., et al. High-performance and site-directed in utero electroporation by a triple-electrode probe. Nature communications 3, 960 (2012).

55. Franklin, K.B.J. & Paxinos, G. The Mouse Brain in Stereotaxic coordinates, (AP, 2008).

56. Heubl, M., et al. GABAA receptor dependent synaptic inhibition rapidly tunes KCC2 activity via the Cl(-)-sensitive WNK1 kinase. Nature communications 8, 1776 (2017).

57. Zhang, J., et al. Modulation of brain cation-Cl(-) cotransport via the SPAK kinase inhibitor ZT-1a. Nature communications 11, 78 (2020).

58. Friedel, P., et al. WNK1-regulated inhibitory phosphorylation of the KCC2 cotransporter maintains the depolarizing action of GABA in immature neurons. Science signaling 8, ra65 (2015).

